# An Integrated Process Analytical Platform for Automated Monitoring of Monoclonal Antibody N-linked Glycosylation

**DOI:** 10.1101/2021.11.14.468439

**Authors:** Aron Gyorgypal, Shishir P.S. Chundawat

## Abstract

The biopharmaceutical industry is transitioning towards adoption of continuous biomanufacturing practices that are often more flexible and efficient than traditional batch processes. Regulatory agencies such as the Food and Drug Administration (FDA) are further urging use of advanced PAT to analyze the design space to increase process knowledge and enable high quality biologics production. Post-translational modification of proteins, such as N-linked glycosylation are often critical quality attributes known to affect biologics safety and efficacy hence requiring close monitoring during manufacturing. Here, we developed an online sequential-injection based PAT system, called N-GLYcanyzer, that can rapidly monitor mAb glycosylation during upstream biomanufacturing. The key innovation includes design of an integrated mAb sampling and derivation system for antibody titer and glycoform analysis in under 2 hours. The N-GLYcanyzer process includes mAb capture, deglycosylation, fluorescent glycan labeling, and glycan enrichment for direct injection and analysis on an integrated high performance liquid chromatography (HPLC) system. Different fluorescent tags and reductants were tested to maximize glycan labeling efficiency under aqueous conditions, while porous graphitized carbon (PGC) was studied for optimum glycan recovery and enrichment. We find that 2-AB labeling of glycans with 2-picoline borane as a reducing agent, using the N-GLYcanyzer workflow, gives higher glycan labeling efficiency under aqueous conditions leading to upwards of a 5-fold increase in fluorescent products intensity. Finally, we showcase how the N-GLYcanyzer platform can be implemented at/on-line to an upstream bioreactor for automated and near real-time glycosylation monitoring of a Trastuzumab biosimilar produced by Chinese Hamster Ovary (CHO) cells.

**Graphical Abstract:** N-GLYcanyzer is an automated PAT toolkit for rapid sample processing for mAb N-linked glycans analysis to enable advanced biologics manufacturing

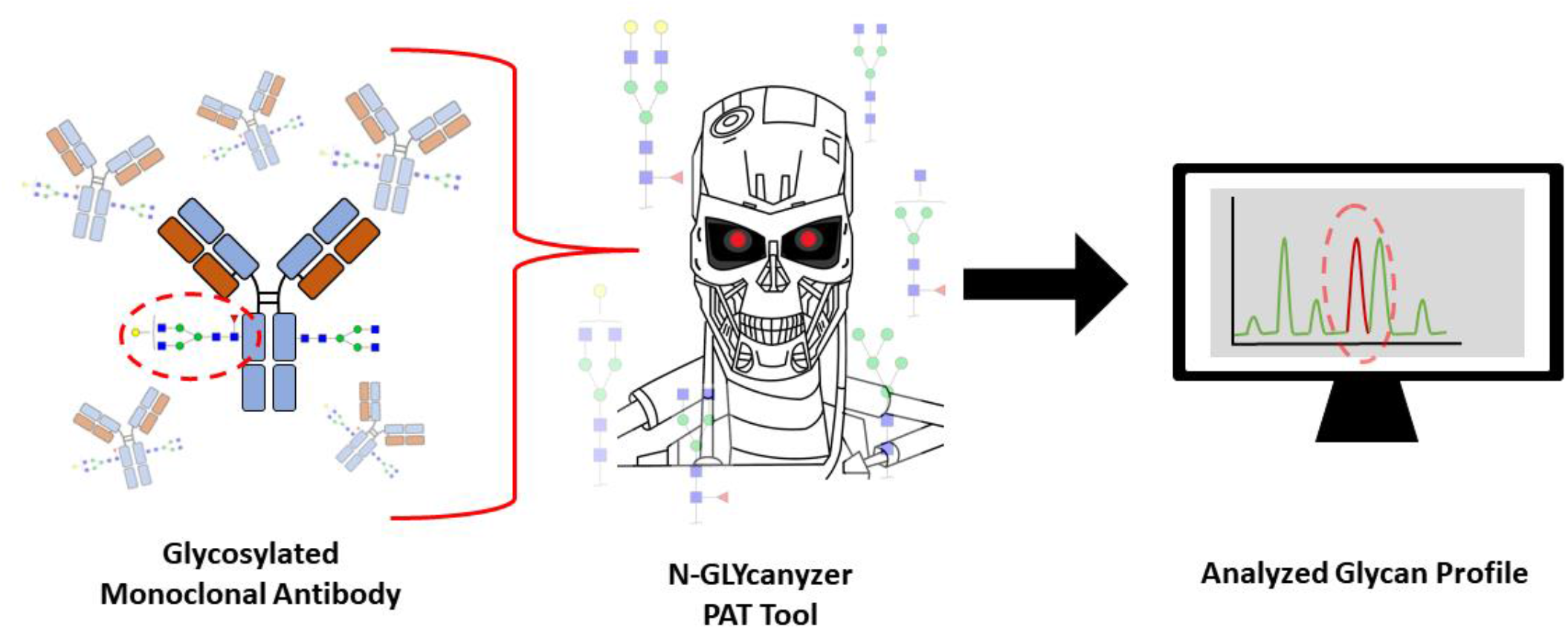

## Introduction

The biopharmaceutical industry is transitioning towards implementation of continuous biomanufacturing approaches for biological drugs manufacturing^1,2^. Continuous bioprocesses can increase production flexibility, simplify process scale-up, improve product quality, reduce process footprint, increase productivity, and reduce production costs^3^. However, the transition from fed-batch to continuous processes for biologics manufacturing is non-trivial and requires contributions from multiple emerging fields such as bioprocess integration, automation, digitization, along with availability of clear regulatory guidelines to facilitate this transition. One key challenge is enabling robust and reliable decision-making during continuous drug production, which relies on real-time monitoring and control of the process^4–6^. Process analytical technology (PAT) can facilitate critical decisions during bioprocessing by monitoring critical quality attributes (CQA’s) that influence the efficacy and safety of the drug substance. The goal of PAT is to ensure consistent quality of the final product by monitoring key CQA’s of the drug product and its intermediates during the entire manufacturing cycle. The Food and Drug Administration (FDA) has endorsed this philosophy and published guidelines for investigation into innovative techniques that can enhance understanding and control of manufacturing processes^7^. The shift towards quality-by-design (QbD) is intended to improve product quality and has been the focus of the biopharmaceutical industry for optimization of batch/fed-batch processes yielding high productivity (e.g., >10 g/L mAb titers)^8^ and further transitioning into advanced/continuous manufacturing processes.

CQA’s can be described as any modification that may alter the characteristics of a drug, such as stability, safety, clearance, or overall efficacy. Different categories of process CQA’s exist such as mAb protein size/aggregation related variants, charge variants, oxidation variants, glycosylation, and other structural-modification variants^9^. Of these CQA’s glycosylation has been of keen interest, especially since specific glycoforms are known to influence the efficacy and safety of biologics^10^. The mAb glycosylation process is heterogeneous and dependent on cell culture and processing conditions, as well as culture media and feed characteristics, which makes process control inherently difficult^11–14^. Current practices in the biopharmaceutical industry only characterize mAb glycosylation often at the very end of the cell culture process because of the time and labor involved in sample preparation for N-linked glycans analysis.

Researchers have therefore explored innovative solutions that can decrease overall processing times associated with sample preparation for glycosylation analysis. Several companies have developed commercial 96-well microplate-based kits for mAb glycosylation analysis using at-line or offline automated liquid handling platforms integrated with HPLC and mass spectrometry (MS), such as the Agilent Instant PC kit chemistry^15^ and the Waters RapiFlour-MS kit^16^. While commercial kits are viable for upstream cell line development and high throughput screening often in offline sample preparation/analysis modes, they are associated with high costs and are often not feasible for real-time monitoring in a biomanufacturing environment. Other techniques may speed up sample preparation at different stages of the overall workflow, such as during enzymatic deglycosylation, chemical labeling, and/or labeled glycan recovery/enrichment. Different glycan tagging methods have been explored, such as the use of aldehyde reactive tags for MS analysis which have enhanced detection sensitivity. Additional instant labeling chemistry based glycan labeling tags such as Instant-PC by Agilent, Rapiflour-MS tag by Waters, and V-tag by Ludger have been developed in recent years^17^. Advanced enrichment methods have also been explored to facilitate labeled glycan recovery. For example, Chu et al. used reactivity driven dye clean-up protocols as an alternative for solid phase extraction (SPE) for glycan enrichment, where they specifically used octanal as liquid-liquid extraction/reaction solvent that showed minimal sample loss but allowed excess 2-AB removal^18^. Alternatively, Wu et al. used bacterial cellulose as an efficient and low cost SPE enrichment matrix for rapid labeled glycan cleanup in under 10 minutes^19^.

Recent innovations have also resulted in PAT tools that can measure mAb glycosylation in near-real-time. For example, Sha et al. designed an at-line sample preparation methodology for labeling N-glycans using a multiplexed magnetic bead-based Protein A purification and Diol solid phases extraction (SPE) enrichment step^20^. Similarly, Tharmalingam et al. have outlined a sequential injection technology based workflow for online mAb sampling and enzymatically released N-glycans preparation^21^. However, one of the major limitations of previous studies has been the lack of an optimized and fully automated workflow that can be readily implemented in a biomanufacturing environment.

Here we developed a sequential injection analysis (SIA) methodology to create a fully automated and optimized online PAT toolkit, called the N-GLYcanyzer, for real-time analysis of mAb N-linked glycans (**Figure 1**). Figure 1 provides a schematic overview of the N-GLYcanyzer PAT workflow integrated upstream to a mammalian cell-based bioreactor producing mAb. The methodology includes sample drawing from a bioreactor using a cell-free sampling probe, affinity based Protein-A chromatography to remove contaminants, concentrate mAb and determine titer, followed by mAb denaturation and deglycosylation using PNGase F, 2-AB fluorescent tag labeling of released N-glycans, and finally PGC enrichment/recovery of labeled glycans before injection onto a hydrophilic interaction chromatography (HILIC) analytical column through an external valve onto an integrated at-line mobile HPLC system equipped with a fluorescence detector. Various labeling chemistries and enrichment steps were studied under SIA compatible aqueous flow conditions to systematically understand how different fluorophore tags and reducing reagents for reducing end glycans labeling influences the overall analytical workflow. We finally validated our online methodology using a generic mAb biologic (Trastuzumab) and compared the N-GLYcanyzer workflow to a classical offline analysis method.

**Figure 1.**
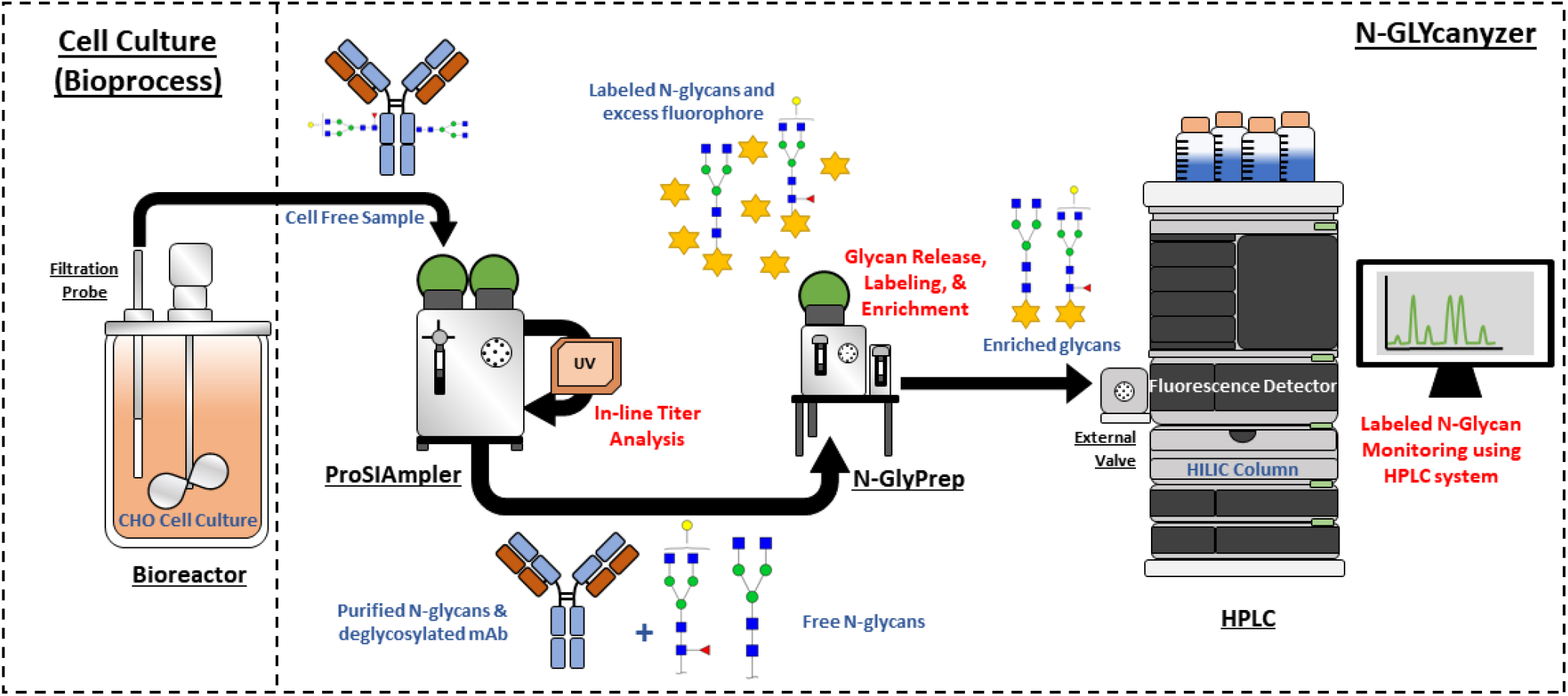
Schematic overview to N-GLYcanyzer PAT workflow system integrated with an upstream bioreactor. Briefly, cell-free culture samples is removed from the bioreactor using a filtration probe to capture mAb using the ProSIAmpler system where protein A chromatography is done to capture mAbs followed by integrated in-line mAb concentration analysis using an inline UV detector. The isolated mAb sample is then sent to the N-GLYprep unit where N-Glycans are released from denatured mAbs using rapid enzymatic digestion. The released N-glycans are fluorescently labeled using a suitable dye/reagents and enriched from the excess unreacted fluorophore dye followed by injection and analysis using an integrated HPLC system equipped with a fluorescence detector.

## Materials and Methods

### Cell Line and Shake Flask Batch Cell Culture

The Chinese Hamster Ovary-glutamine synthetase (CHO-GS) engineered cell line for production of Trastuzumab was kindly provided by GenScript Biotech Corporation (Piscataway, NJ). One ampule of cells from a working cell bank was thawed in CD-CHO basal medium (Thermo Fisher Scientific, Waltham, MA), along with 0.5 % anti-clumping agent (Thermo Fisher Scientific, Waltham, MA) and 25 μM methionine sulfoximine (MSX) in a 125 mL unbaffled shake flask (VWR, Radnor, PA, USA) with 40 mL working volume. The cells were grown at 37 °C, 130 RPM, and 5% CO2 in a New Brunswick S41i CO2 incubator shaker (New Brunswick Eppendorf, Hamburg, Germany) for 4 days and passaged twice to 0.5×10^6^ cells/mL in a 250 mL and 500 mL shake flask, respectively, and grown for 4 days prior to bioreactor inoculation.

### Bioreactor Fed-Batch Cell Culture

The bioreactor experiments were conducted in a 3L glass bioreactor (Bioflo 320, Eppendorf, Enfield, CT) with a working volume of 1.75 L. The bioreactor temperature and pH control were initiated prior to seed cells inoculation. The set points for dissolved oxygen (DO) concentration, pH, and temperature were 64%, 7.1, and 37 °C, respectively. The pH was controlled by adding either 0.5 M sodium bicarbonate (Sigma Aldrich, St. Louis, MO) or sparging of CO_2_. The bioreactors were sampled daily to analyze various cell culture parameters (e.g., glucose, lactate, glutamate, glutamine, Na^+^, K^+^, Ca^2+^, NH_3_), along with media osmolarity, measured using the BioProfile FLEX2 Analyzer (Nova Biomedical, Waltham, MA). Total cell concentration and viability was measured using the trypan blue exclusion method using trypan blue solution (Sigma Aldrich, St. Louis, MO). The glucose concentration was measured and maintained between 3-5 g/L by supplementation of 500 g/L glucose stock solution. The cell culture in reactor was further fed Dynamis medium (Gibco, Thermo Fisher Scientific Inc., Waltham, MA), as a 5% v/v bolus addition of feed added every other day after day 4 until termination of cell culture on day 14.

### Offline Dextran Oligomers and mAb N-linked Glycans Sample Preparation/Analysis

Lyophilized dextran ladder standard (Sigma-Aldrich, St. Louis, MO) was reconstituted into Milli-Q water (Millipore Corporation, Billerica, MA) to a concentration of 2 g/L. Next, 50 µL of dextran solution was mixed with 50 µL of labeling solution (85% Dimethyl Sulfoxide (DMSO) 15% glacial acetic acid) consisting of either 48 g/L 2-aminobenzamide (2-AB) or anthranilic acid (2-AA) with either 0.5M sodium cyanoborohydride or 2-Picoline Borane (2-PB). The mixture was incubated at 65 °C for 2 hours to allow for efficient labeling. Next, the sample was cooled and diluted 5-fold into HPLC grade water and cleaned using a PGC column (Hypersep Hypercarb, Thermo Fischer Scientific, Waltham, MA). Samples loaded onto PGC column were washed with aqueous based solution of 5% Acetonitrile (ACN) + 0.1% trifluoroacetic acid (TFA) and then eluted in 90% ACN + 0.1% TFA based aqueous solution. All residual liquid was removed from the eluted labeled glycan sample using a Savant speed vac concentrator centrifuge (Savant Instruments, Farmingdale, NY) and then reconstituted in 70% Acetonitrile for HPLC analysis using an Advanced-Bio Glycan Mapping Column Agilent Technologies, Santa Clara, CA) on an Agilent 1260 Bioinert HPLC system equipped with a fluorescence detector together on a mobile cart.

The same methodology was followed for labeling N-glycans released from monoclonal antibody. Briefly, 100 µg of Protein A purified and neutralized mAb (45 µL) was mixed with 2.5 µL denaturation solution to a final concentration of 40mM Dithiothreitol (DTT) and 0.1% sodium dodecyl sulfate (SDS), heated to 95 °C for 5 minutes, and cooled. Non-ionic surfactant NP-40 (Sigma-Aldrich, St. Louis, MO) was added to a final concentration of 0.5% along with 2 µL of PNGase F Ultra (Agilent Technologies, Santa Clara, CA) and incubated for 1 hour at 37 °C before subsequent 2-AB/2-AA labelling, enrichment, and HPLC analysis as described above.

### Online mAb Sampling, Titer Determination, and N-linked Glycan Processing Workflow

A cell-free culture sample was removed from the bioreactor culture using a 0.22 µm filtration probe (Trace Analytics, Braunschweig, Germany) and pumped into the N-GLYcanyzer workflow unit. This unit overall consists of a custom-built and fully integrated instrument consisting of a ProSIAmpler device (FIAlab Instruments Inc., Seattle, WA) and two syringe pumps and one 9 port valve we have named the N-GlyPrep system, programmed in PYTHON using SIASoft v1.1.7 (FIAlab Instruments Inc, Seattle, WA) as schematically outlined in **Figure 2**. Briefly, the cell culture sample was injected onto a Protein A (ProA) column packed with mAbselect SuRe protein A resin (Cytiva, Marlborough, MA), and washed with 20 mM sodium phosphate buffer pH 7.0 and then eluted with 0.1% formic acid pH 2.5. Titer was determined using a miniaturized in-line UV spectrometer (Ocean Optics, Dunedin, FL) integrated downstream of the ProA column. Eluent was adjusted to neutral pH with 1 M Tris pH 9.0 buffer. The eluted mAb was then mixed with denaturation solution to a final concentration of 0.1% SDS and 40mM DTT, and then incubated at 95 °C for 5 minutes. The sample was allowed to cool to room temperature prior to addition of NP-40 to a final concentration of 0.5%. Next, 2 µL of PNGase F was added and the mixture was incubated at 37 °C for 5 mins (rapid digestion) and up to 1 hour (standard digestion). The mixture of glycans was combined with the 2-AB labeling solution (i.e., 48 mg/mL 2-AB, 1 M 2-PB in 70:30 DMSO, Glacial Acetic Acid) and left to react in a reaction coil at 65 °C for 2 hours. After cooling, the sample was diluted 5-fold into HPLC grade water and injected onto a self-packed PGC column using Hypercarb material. The loaded sample was cleaned from background impurities using 5% ACN + 0.1% TFA wash solution then eluted in 90% ACN + 0.1% TFA based aqueous solution. The eluent was then directly transferred via an Agilent 1290 external switching valve inline to the HPLC system, and automatically prompted by SIAsoft the HPLC for running the HILIC analysis. A detailed workflow is shown in Figure 2.

**Figure 2.**
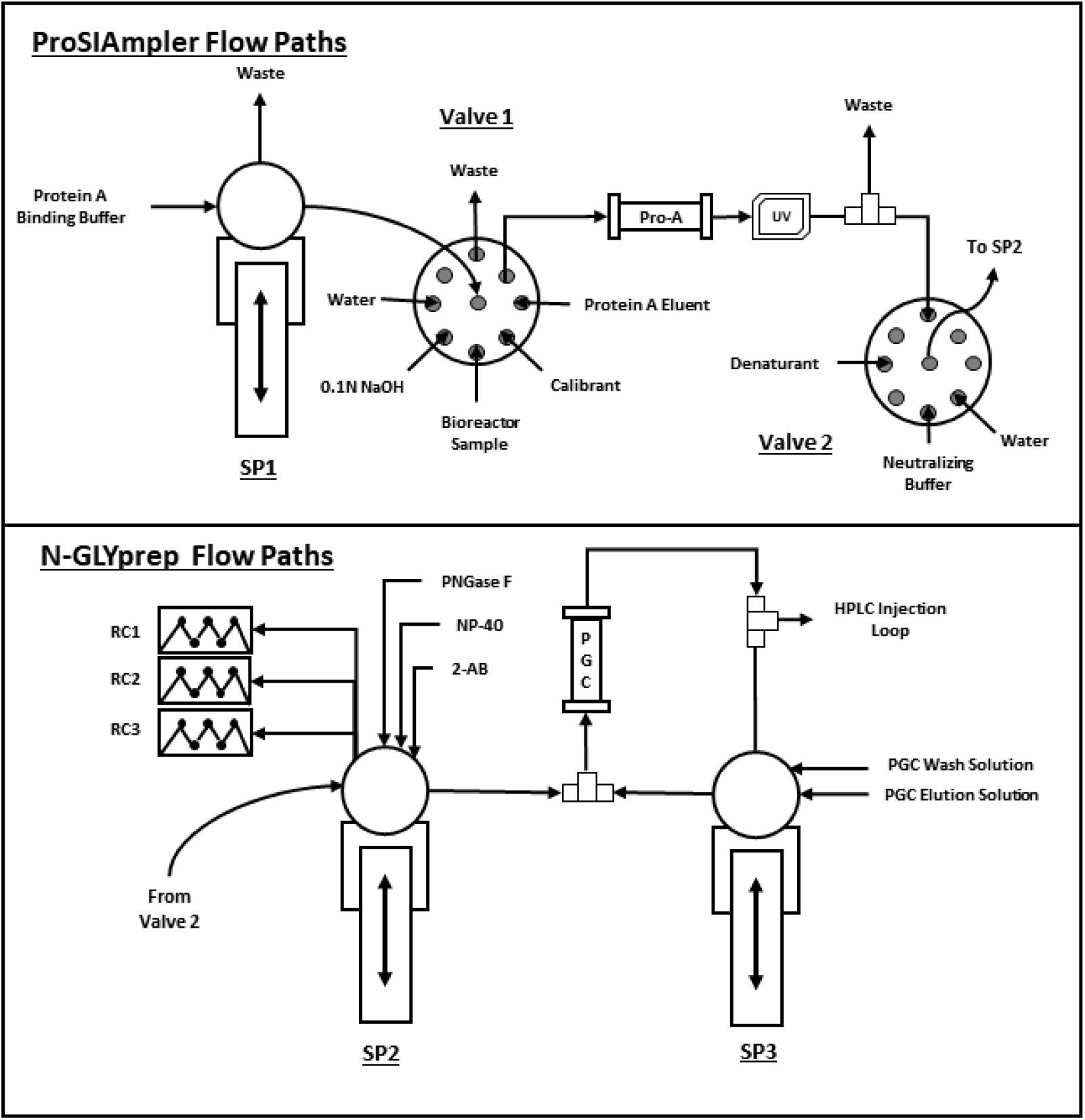
Detailed N-GLYcanyzer fluidics process flow diagram for real-time mAb titer and glycosylation monitoring. The figure above shows the sample uptake via Valve 1 on the ProSIAmpler unit where the cell free sample is loaded onto a Protein A column, cleaned using a ProA column binding buffer using SP1 (Syringe Pump 1) and then eluted using Eluent mobile phase via Valve 1. The eluted mAb sample is then pumped through an inline UV spectrophotometer to measure UV=280nm absorbance to estimate mAb titer based on a pre-determined calibration curve. The sample is then pumped through valve 2 on the N-GLYprep unit along with introduction of a neutralizing buffer to increase the pH and add a denaturant followed by mixing using SP2 (Syringe Pump 2) and then injected into reaction coil 1 (RC1) set at 90 °C for 10 minutes. Samples are then removed and NP40 is added, followed by cooling to room temperature, and then PNGase F enzyme is added and reaction mixture is sent to reaction coil 2 (RC2) set at 37 °C for 5 mins to 1 hour. The samples mixture is afterwards mixed with 2-AB reagent solution and injected into reaction coil 3 (RC3) set at 65 °C for 2 hours. The sample is then removed, cooled and diluted with water for injection via valve 2 to the PGC column. Samples are washed with PGC wash solution and eluted with elution solution, via syringe pump 3 (SP3). Eluted samples are mixed at SP3 and then pumped into the injection loop on the HPLC.

## Results and Discussions

### Accurate At-line mAb Sampling & Titer Analysis Enabled using ProSIAmpler

Efficient at-line mAb sampling from the bioreactor and accurate titer determination was critical to optimize the downstream N-glycan release protocols and labeling chemistry. Monoclonal antibody titer was determined against a calibration curve using a custom built ProSIAmpler sample draw system integrated with the bioreactor. The 7-point calibration curve was made using an in-house generated mAb standard by preparing samples between 0 – 2.0 g/L of mAb in 100 µL of protein A eluent solution. **Figure 3**A shows the calibration curve generated for the protein A column, where the standards show a strong linear correlation between the integrated 280nm absorbance peak areas and mAb concentration for n=3 replicate calibration standard injections into the ProSIAmpler UV flowcell. Figure 3B shows the UV elution profile for the 1.5 g/L mAb standard injection which also showed strong reproducibility between replicate injections.

**Figure 3.**
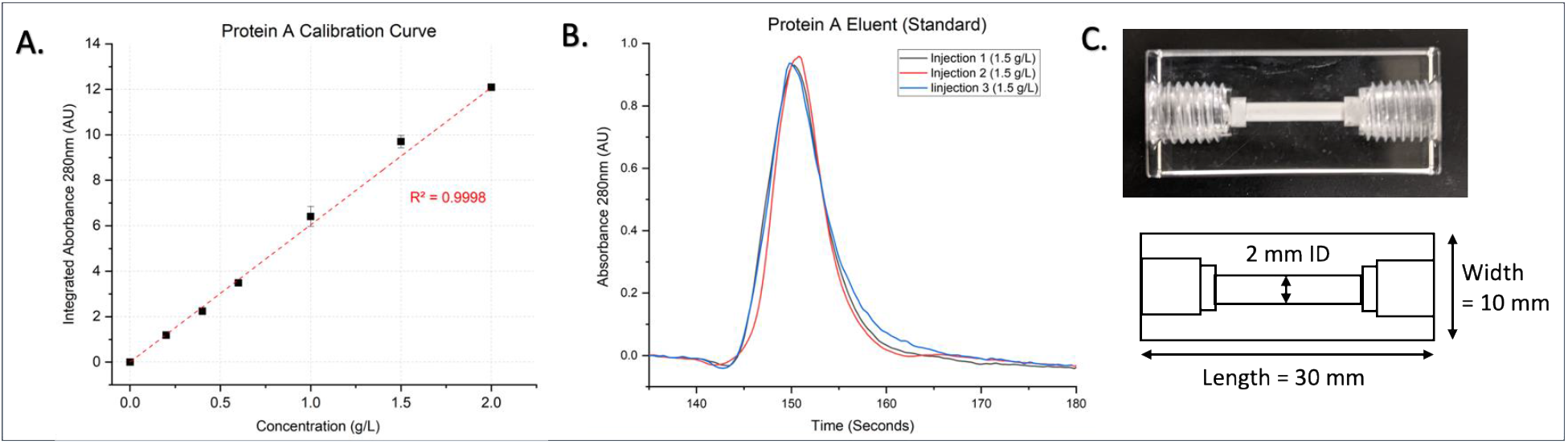
ProSIAmpler unit allows accurate mAb recovery and protein titer determination. Here, (3A) shows the 7 points-based calibration curve made by injection of 0 to 200 µ g of mAb at a constant 100 µL injection volume that showed a linear correlation with r^2^ = 0.9998. (3B) shows a representative chromatogram obtained from the inline UV spectrophotometer for three equivalent injections of a 1.5 g/L mAb solutions. We see a very consistent residence time on the pre-packed ProA column. (3C) shows the self-packed protein A column dimensions and representative picture of an actual column.

The dimension of the packed ProA column can be seen in figure 3C. The self-packed protein A column was made from Poly(methyl 2-methylpropenoate) material, or plexiglass, which is known to have strong resistance towards dilute acids. The protein A column was sealed on both ends using a porous cellulose frit suitable for chromatography. Longer column widths and wider column diameters were also tested but it caused peak distortion issues due to increase in residence time and unwanted mixing within the column. The column was also tested for low and high concentration ranges of protein binding between 0 to 720 µg of mAb loading. The column showed strong correlation up to 200 µg of mAb binding but showed slight deterioration in binding recovery with higher loadings as seen in SI Appendix Fig. S1. It should be noted that using an eluent with the addition of ionic strength modifier such as phosphate buffer at a lower pH could help to increase elution performance. However, we decided against the use of such modifiers to decrease the total salt concentration within our workflow that could interfere with downstream sample processing. Similar mAb binding experiments have been reported on a similar ProSIAmpler system by Liu et al^22^ that also gave very similar results to when using a commercially available and pre-packed protein A columns.

### Optimization of Fluorophore Labeling under Aqueous versus Non-aqueous Reaction Conditions

Two fluorescent glycan-tagging reagents were used to test the labeling chemistry efficiency under aqueous/non-aqueous conditions, using multiple reductants to complete the Schiff base reduction reaction, and compare fluorophore signal-to-noise ratio using a model dextran ladder standard. Namely, 2-AB and 2-AA were compared, where 2-AB differs from 2-AA as it contains an amide group rather than a carboxylic acid moiety and used along with 2-Cyanoborohydride and 2-Picoline Borane as reductants. A similar study was done by Kozak et al comparing 2-AB with Procainamide as the fluorescent label for glycans^23^. The use of a toxic reducing agent such as sodium cyanoborohydride presents concerns for the buildup of hydrogen cyanide as a side reaction during Schiff-base formation, making it challenging to establish a real-time PAT toolkit used at-line during biomanufacturing operations. 2-PB has been documented as a non-toxic alternative for reducing the Schiff base of conjugated oligosaccharides to a fluorescent tag^24^. Hence, we were interested to compare the efficiency of using 2-PB as an alternative reducing agent to sodium cyanoborohydride for implementation in our online PAT methodology.

Traditional methods for fluorescent dye labeling of reducing sugars are mostly reported under non-aqueous conditions using glacial acetic acid and DMSO as the reaction solvents. Before conjugation of the fluorescent dye, samples need to be either lyophilized or dried offline in a speed-vac to remove any residual traces of aqueous solvents. This is because water is a byproduct of the Schiff base formation reaction during generation of the imine intermediate^25^. The accumulation of water in the reaction solution that is highly aqueous could potentially inhibit the formation of the Schiff base and the subsequent reduction step to reduce fluorophore labeling efficiency. However, the online automated sample preparation workflows ideally cannot be designed to remove residual aqueous solvents from the sample, as often done during offline manual sample preparation workflows for N-glycan labeling. Since there was limited information available about the labeling efficiency of glycans under aqueous versus non-aqueous conditions, we determined the reaction efficiencies for 2-AB/2-AA under different labeling conditions. Similar preliminary experiments were reported by Ruhaak et al. to test if 2-PB would be a suitable alternative to sodium cyanoborohydride^24^. However, they did not test the use of 2-AA under aqueous conditions, which was further investigated in our work along with providing a detailed quantitative analysis of all detected product profiles.

For the samples labeled under aqueous conditions, we first tested a model dextran ladder standard (100 µg) dissolved in 50 µL of water followed by addition of 50 µL of labeling mixtures (i.e., 1 M 2-PB or sodium cyanoborohydride along with 48 g/L 2-AB or 2-AA dye dissolved in 85:15 DMSO:Glacial Acetic Acid solution) that was mixed and heated at 65 °C for 2 hours. For the samples labeled under non-aqueous conditions an equivalent amount of dextran, but in the absence of water, was reconstituted into a labeling solution (i.e., 1M 2-PB or sodium cyanoborohydride with 48 g/L 2-AB or 2AA in 85:15 DMSO, Glacial Acetic Acid), mixed and heated at 65C for 2 hours. This allowed us to keep the molar concentrations consistent between the aqueous versus non-aqueous reaction conditions. Control samples were prepared in the absence of any reductants to understand the glycan labeling efficiencies without reducing the formed Schiff base. Results for these sets of experiments are reported in **Figure 4A**. For the dextran ladder, we have only shown the results for the 10-mer oligomer of glucose (i.e., maltodecaose) as an analog to similarly sized N-glycans expected to be released from a glycosylated monoclonal antibody. However, as the degree of polymerization for mAb N-glycans range can vary drastically depending on glycan maturation, additional results for other size oligomers of glucose and their labelling efficiencies are reported in the supplementary information (Fig. S2). All values reported are normalized to the highest fluorescent signal at the optimal excitation and emission wavelengths of the fluorophores (i.e., 260 nm excitation and 430 nm emission for both 2-AB and 2-AA).

**Figure 4.**
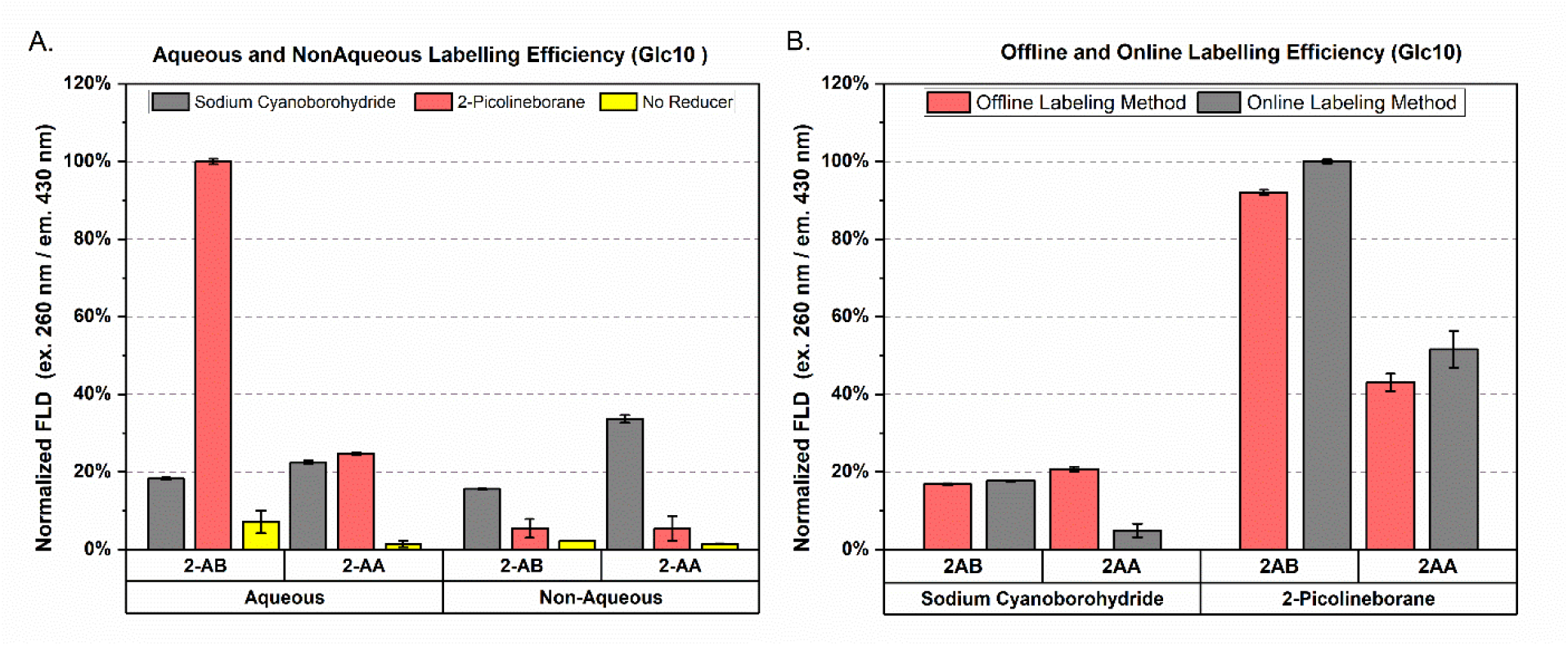
2-Picoline borane outperforms sodium cyanoborohydride as a reducing agent under aqueous conditions for 2-AB fluorophore labeling of dextran. Here, (4A) shows the labeling efficiency of the 10-mer of glucose (i.e., maltodecaose) in the dextran ladder. (4B) shows no biases between the online versus offline labeling methods when 2-picoline borane was used as the reductant for both 2-AB/2-AA as fluorophore tags.

For the aqueous conditions, surprisingly we noted that reduction efficiency of the Schiff base with 2-PB was significantly higher than that seen with sodium cyanoborohydride for both 2-AA and 2-AB. This finding is also in agreement with a similar investigation reported on plant cell wall mono and disaccharides fluorophore labeling by Fang et al^26^ which showed that 2AB labeled sugars yielded higher efficiencies when 2-PB was used as the reductant over sodium cyanoborohydride. Aqueous 2-AB labeling with 2-PB showed the highest fluorescent intensity and was used to normalize results from all other conditions reported. 2-AA labelling with 2-PB followed with the second highest fluorescence intensity (23% normalized), while the use of cyanoborohydride as the reductant under the aqueous conditions gave much lower intensities for both 2-AB (18% normalized) and 2-AA (22% normalized) labeling. These results differ from earlier reports when comparing the fluorescent tag labeling sensitivity between 2AA and 2-AB, however the prior work did not use 2-PB as the reducing agent which likely contributes to the overall labeling efficiency^27,28^. We hypothesize that addition of water could increase the efficiency of the reduction reaction as 2-PB is miscible with water, which could enhance the rate of Schiff base formation and its reduction. For non-aqueous conditions, we noted that sodium cyanoborohydride clearly outperformed 2-PB as a reductant based on labeling efficiency. This suggests that the increased acidity of 2-AA over 2-AB under non-aqueous conditions could enhance the performance of 2-PB as a reductant, as increased acidity would favor the open chair pyranose ring conformation of the reducing end moiety leading to an increase in fluorescent dye conjugation to the oligosaccharide although the non-aqueous results for 2AB + 2 PB and 2-AA + 2-PB both showed comparable labeling efficiencies.

Samples were next labeled under aqueous conditions using our automated online N-GLYcanyzer workflow and compared to the manual offline method. All samples and reagent solutions were prepared as described above for aqueous sample labeling conditions. Offline analysis samples were incubated on a heat-block at 65 °C in the dark for 2 hours, while the online samples were mixed aspirated using the online N-GLYcanyzer unit, mixed, and incubated into a heating coil at 65 °C for 2 hours. All samples were then cleaned using a PGC SPE cartridge as described under the offline sample preparation protocol earlier. The relative labeling efficiencies are shown in Figure 4B. The online methodology shows marginally higher labeling efficiencies versus the offline method in all cases. This can be largely attributed to more homogeneous heating of the reactant mixture in the N-GLYprep heating coil versus microfuge tubes in the heat block.

With these results, we decided to use 2-AB as the preferred labeling dye for our online mAb N-glycan preparation/labeling method and use 2-PB as the reductant for completion of the subsequent PAT method objectives.

### Labeled Glycan Enrichment & Purification

Labeled glycan enrichment, also known as labeled glycan cleaning/purification, is an essential step before HPLC analysis to remove excess labeling dye and other background reagents from the sample that could potentially interfere with the chromatography. Since 2-PB showed a better labeling efficiency versus the traditional sodium cyanoborohydride reductant reported the literature, we were interested to understand how both reductants may potentially interact with the PGC material to impact labeled glycan enrichment and recovery^26,29^. Labeled glycans have been reported to have sufficient retention on PGC based chromatographic materials^30–32^, and has also been used for online workflows prior^33^. Therefore, we primarily focused on use of PGC materials for labeled glycan enrichment but also explored other chromatographic materials that is not reported here (e.g., HILIC). To optimize the enrichment step, the mass of the labeled glycan sample reduced with 2-PB and recovered was calculated as a function of PGC bed weight/volume used for the enrichment/cleanup step. The PGC bed weight was varied between 10 to 100 mg (dry weight basis) and optimized for the online workflow once again using a dextran ladder as a glycan analog. The use of the highest recovery data point (i.e., 100 mg PGC with 2-PB as the reductant) was used to normalize all values for comparison at the varying PGC bed weights. First, dextran ladder standard based samples were prepared as previously described for aqueous labeling. Next, 100 µg of dextran was labeled with 2AB using either sodium cyanoborohydride or 2-PB as the reductant at the equivalent molar concentrations. Samples were then diluted fivefold with water to a volume of 500 µL before glycan enrichment using a PGC column of varying bed weights.

**Figure 5A** shows the labeled sample recoveries of 2-AB labeled glucose ladder 8-mer, 10-mer and 12-mer using either sodium cyanoborohydride or 2-PB as the reducing agent. Surprisingly, sodium cyanoborohydride reduced glycans dextran showed a higher rate of PGC recovery versus 2-PB on average before the 100 mg loading point. Details on recoveries of other oligomers of glucose from the labeled dextran ladder standard can be found in the supplementary information (Fig. S3). These results suggest that if using sodium cyanoborohydride as a reductant a lower bed weight of PGC can be used but with only modest recoveries than that seen with 2-PB. This can be mitigated as the bed weight is increased to increase the number of available PGC binding sites for labeled glycans adsorption. This explains the linear trend in increased glycan recovery of 2-PB reduced samples as the PGC bed-weight was increased. Alternatively, the molarity of 2-PB used in solution may also be decreased if it does not compromise the efficiency of the Schiff base formation and reduction. The relative recovery was close to 100% under the optimal PGC bed weight (100 mg) for the 8-mer, 10-mer, as well as for 12-mer from the labeled dextran ladder, when using 2-PB as the reducing agent, along with robust and reproducible results in sample recoveries. Recovery values for overall labeled glycan recoveries show a similar trend for all dextran oligomers of increasing molecular weights at the optimum PGC bed weight of 100 mg. However, at 2.5-lower lower PGC bed weights to the optimum amount (40 mg vs. 100 mg PGC) glycan recovery was noted to be higher for the larger dextran (e.g., 8-to-12-mer) versus smaller dextran (e.g., 4-to7-mers) oligosaccharides. Similar increased recovery and tighter adsorption affinity for higher molecular weight glycans to PGC has been often reported in the literature.

**Figure 5.**
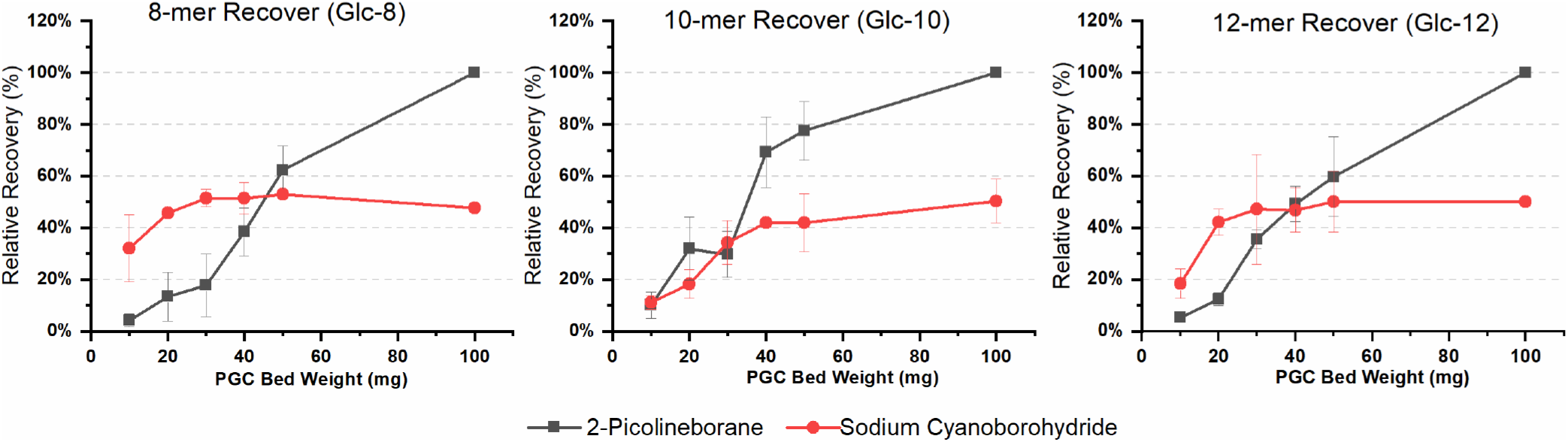
Oligosaccharide recovery for labeled dextran ladder depends on reducing agent and PGC amount. All values are normalized to the highest recovery based on the specific 8-mer, 10-mer and 12-mer oligomers of glucose (i.e., maltooctaose, maltodecaose, and maltododecaose, respectively). An increasing trend in total recovery can be seen with all oligomers when 2-picoline borane is used as a reductant while the use of sodium cyanoborohydride seemed to saturate quickly with increasing PGC bed weight/volume.

The redox state of PGC has been previously demonstrated in the literature to impact the retention of polar analytes like 2AB-labeled malto-oligosaccharides and N-linked glycans^34^. PGC surface oxidation can lower glycan adsorption enthalpy/entropy values and is thought to result in increased interactions between the glycan hydroxyl groups and the oxidized PGC surface. Tighter binding of polar analytes to an oxidized PGC surface could reduce the degrees of freedom of the adsorbed molecules which results in lower reported adsorption entropies. Based on these reports, reduction of the PGC surface should result in weaker binding of polar analytes and this could partly explain why lower glycan enrichment/recovery was seen for all reductant containing samples shown in Figure 5. However, surprisingly there seemed to be weaker binding affinity of most 2-AB labeled malto-oligosaccharides to the reduced PGC surface in the presence of 2-PB versus cyanoborohydride. We hypothesize 2-PB has a competitive binding interaction on PGC, possibly because of its pyridinium ring structure that interacts to the graphitic carbon interface via hydrophobic interactions (e.g., CH–π interactions), that additionally decreases reduced PGC surface binding sites occupancy of labeled glycans and results in subsequent loss in recovery. While it was outside of the scope of work reported here, it might be possible to also fine-tune the interplay of enthalpic and entropic contributions to the overall adsorption phenomena using temperature modulation instead of varying total PGC binding sites to maximize the glycan enrichment/separation process in presence of 2-PB reducing agent.

### Online N-Glycan Enzymatic Release, 2-AB Fluorophore Labeling, and PGC Enrichment using N-GLYprep

The online workflow on the N-Glyprep unit mirrored the offline sample preparation scheme, however all volumes were doubled for the analysis to mitigate loss of sample from dead-volumes associated with the system. For example, the off-line sample preparation used a denaturation and deglycosylation reaction volume of about 50 µL while the volumes of the N-Glyprep unit was doubled to 100 µL. After deglycosylation, the sample was homogenized within the syringe (SP1) and half the sample was taken for 2-AB labelling while the second half was moved to waste. The deglycosylation sample was mixed in equal volumes of the labeling mixture and reacted for 2 hours as stated, but these steps can be readily optimized to be run at shorter reaction times. The 100 µL mixture was next diluted 5 fold into Milli-Q water and after PGC column equilibration was complete the sample was added to the column and washed with 5% ACN + 0.1% TFA, and finally the enriched sample was eluted in 90% ACN + 0.1% TFA.

### Integrated N-GLYcanyzer and HPLC Chromatography Analysis Workflow

After labeled sample enrichment, the sample was eluted from the PGC column, homogenized, and aspirated into a 5 µL injection loop on an external valve on the mobile HPLC. Once the sample was placed in the injection loop, the HPLC was triggered by an external start from the SIAsoft software prompting the HPLC run to initiate. After the first 2 minutes of HPLC run initiation, the external valve was switched in line with the column, and the sample was injected into the pre-equilibrated analytical HILIC column for chromatographic separation/detection of the 2-AB labeled glycan analytes. Representative chromatograms comparing the manual offline (as control) and automated online sample preparation workflows can be seen in **Figure 6**. Figure 6A shows the comparison of the dextran sample ladder analyzed on the HPLC using the automated online and offline workflows. Figure 6B shows the HPLC chromatogram from a N-linked glycans released from model mAb (Trastuzumab) after 2-AB labeling and enrichment. Representative chromatograms for online/offline sample analysis workflows are shown here to compare the relative abundances and retention times for each mAb glycoform often reported for Trastuzumab. Results show no bias introduced from the automated online sample preparation workflow (versus the conventional manual offline workflow) and have very similar retention times with minimal standard deviations, suggesting high analytical reproducibility of our developed methodology.

**Figure 6.**
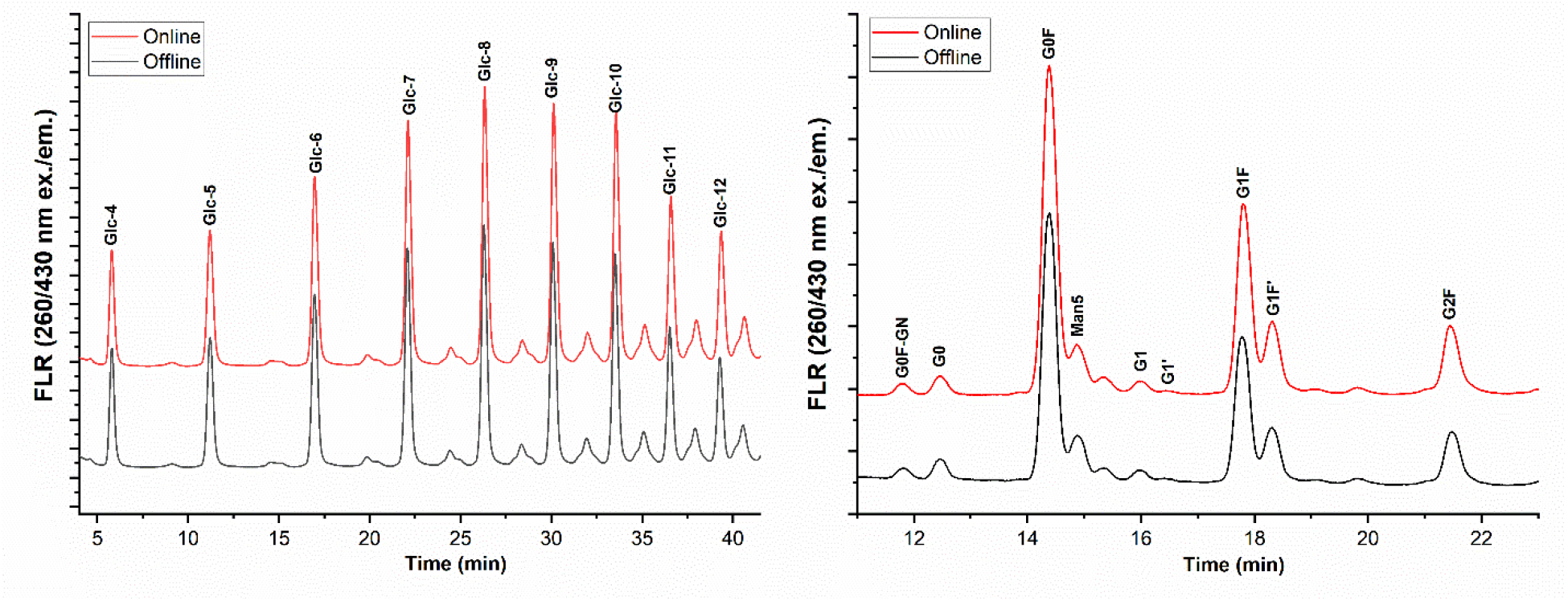
N-GLYcanyzer PAT workflow gives comparable glycan labeling and recovery versus classical offline workflow. A representative chromatogram for the dextran ladder can be seen in figure 6A while mAb N-linked glycan glycoforms from Trastuzumab mAb is shown in 6B. In both cases samples were prepared using the automated N-GLYcanyzer online and manual offline methodologies. Chromatograms show very similar relative fluorescence signals and retention times. Detailed relative abundances and retention times are calculated and reported in table 1.

PGC based sample enrichment was decided over the use of HILIC based materials to mitigate the risks associated with sample solvent mismatch. For example, having a high-water content after sample enrichment using HILIC based materials may cause issues with sample retention on the analytical column as water is a strong elution solvent for later HPLC-HILIC analysis. However, eluting the sample off the PGC column before injection using 90% acetonitrile and 0.1% TFA did not seem to cause any issues with downstream analytical HILIC chromatography on the HPLC. It is assumed that the mobile phase in the HPLC being 80% acetonitrile and 20% 100 mM ammonium formate pH 4.4 diluted the water content of the injected sample sufficiently enough to mitigate potential sample solvent & mobile phase mismatch. Our preliminary experiments also explored eluting samples off the PGC enrichment column using a higher acetonitrile concentration, such as 100% acetonitrile with 0.1% TFA. However, severe eluting peak distortion was noticed during the HPLC analysis when using this injection solvent with higher acetonitrile composition versus that of the HILIC column equilibration mobile phase composition. Using an eluent additive such as ammonium formate with acetonitrile may also increase the elution strength off the column leading to an increase in the chromatogram signal. Additional optimization of the eluent solvent for the PGC enrichment column and the mobile phase for the analytical HPLC-HILIC column are indeed possible, particularly including a HILIC guard column. This could further improve robustness of the sample preloading onto the HILIC column and potentially minimize drift in retention times of analytes over multiple injections over several days/weeks when using the N-GLYcanyzer-HPLC PAT workflow to continuously monitor samples at-line from an upstream bioreactor.

**Table 1:**
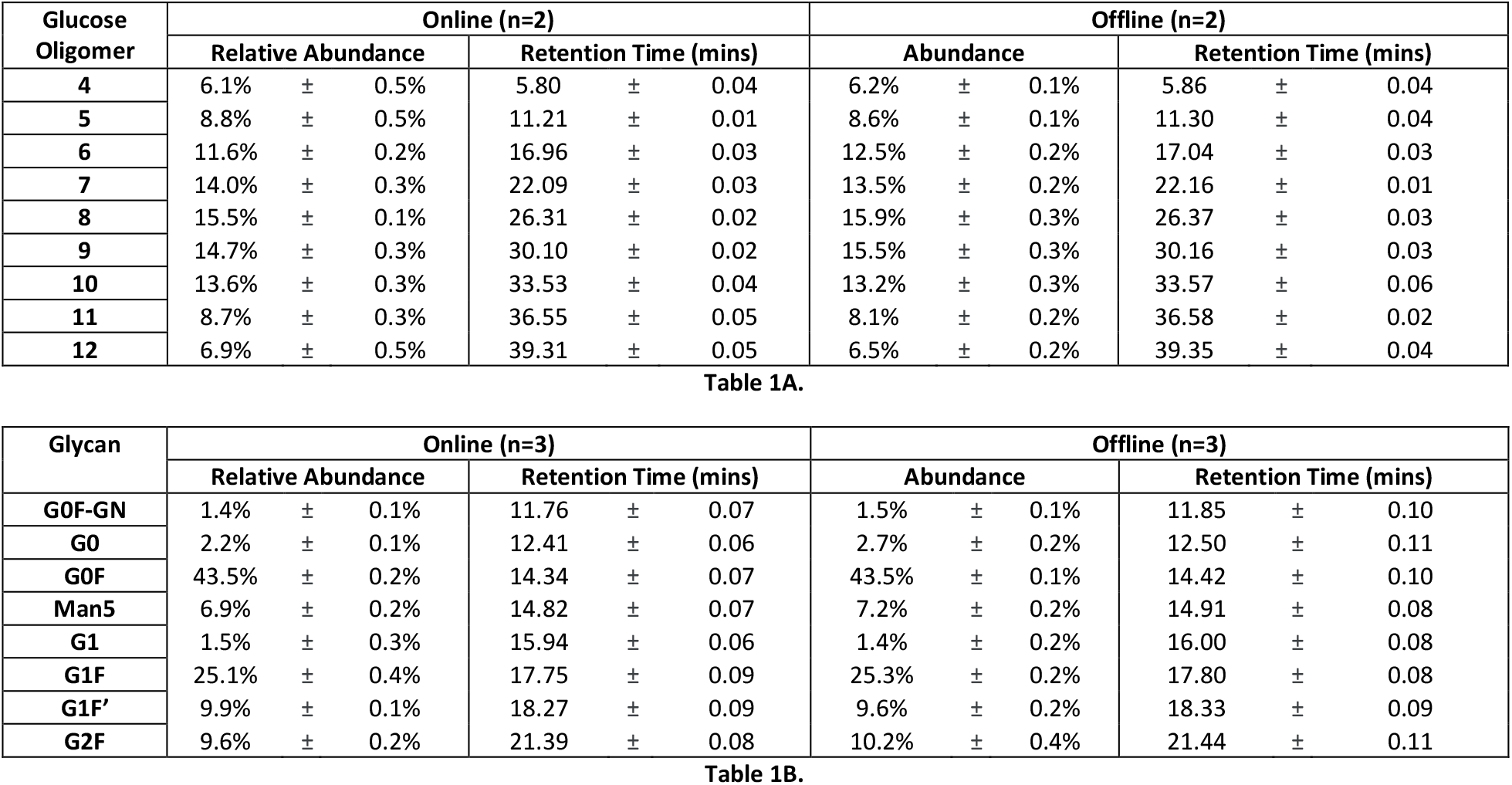
Integrated chromatography peaks for dextran ladder (1A) and mAb N-glycan (1B) samples are shown here. Both results show no significant biases between the online and offline methodology for technical replicates (total number is shown in parentheses). For the dextran oligomers, the standard deviations seemed marginally higher versus the offline method however the differences were not statistically significant.

### Real-Time mAb Glycosylation and Titer Monitoring using N-GLYcanyzer PAT toolkit for CHO Cell Bioreactor Culture

Finally, to demonstrate the utility of our methodology for glycan analysis, we implemented our N-GLYcanyzer PAT toolkit to draw mAb sample and purify using a ProA column followed by N-linked glycan release, labeling, and enrichment for fully automated analysis by an integrated mobile HPLC system. Real-time measurements were performed in a fully autonomous manner to monitor mAb titer from CHO cell culture samples from a fed-batch bioreactor run ranging from day 0 through day 14 as shown in **Figure 7A**. Offline measurements using the standard HPLC based method were compared against the real-time N-glycan measurements, with all analyses run in at-least duplicates. Both offline and online based analysis showed similar precision and gave comparable glycoform abundance results for mAb glycosylation over the duration of the CHO cell culture.

**Figure 7.**
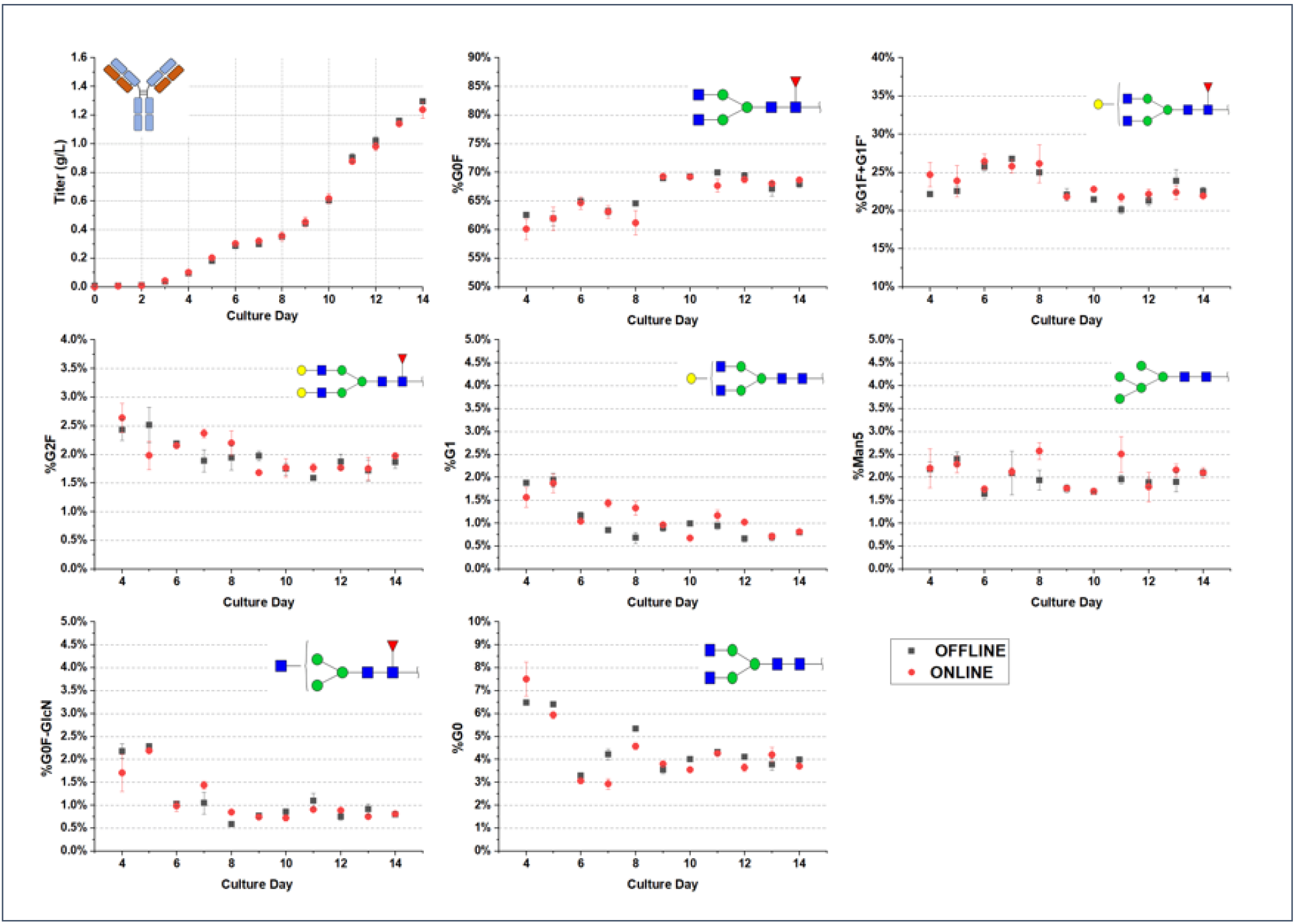
CHO cell culture mAb glycosylation was monitored using the automated N-GLYcanyzer PAT workflow. Online (red filled circles) and offline (black filled squares) analysis of the mAb titer and its major glycoforms are shown here. Titer between the online and offline methods gives very similar and reproducible values. Similarly, comparable results are observed for various glycoforms such as G0F, G1F+G1F’, G2F, G1, Man5, G0F-GlcN, and G0 using both offline and online methods. All values shown are normalized as relative abundances for each glycoform at each respective day. Error bars indicate one standard deviation for reported mean values from replicate analysis.

For released N-glycan analysis, 7 major glycoforms expected for our IgG antibody Trastuzumab were detected using the online methodology and validated against the standard offline analytical workflow. We determined that the standard offline analytical method results mirrored the online method closely. Briefly, an increase in certain glycoforms such G0F (figure 7B) was seen during the progression of the CHO cell culture while a clear decrease in total galactosylation based glycoforms such as G1F+G1F’, G2F and G1 (i.e., Figure7C, 7D and 7E, respectively) was clearly seen using both the online and offline based analytical workflows. Similar trends in change in galactosylation levels for mAbs have been reported previously for several mAb producing CHO cell lines^35^.

## Conclusion

Our study has established and validated a fully integrated and automated methodology for online mAb titer and N-glycosylation analysis using an integrated flow injection analysis system equipped with an inline UV spectrophotometer for real time mAb titer analysis, N-linked glycan release, fluorophore dye labeling of N-glycans, labeled N-glycans enrichment using mini-PGC column optimized under aqueous flow through conditions, and finally downstream HPLC system for fluorescently labeled N-glycan analysis. The proposed system was shown to enable in-/on-/at-line upstream bioprocess monitoring of mAb titer within minutes of drawn sample analysis and ability to monitor glycosylation within 1-3 hours of sample draw continuously over a 14-day CHO cell culture producing a glycosylated mAb biologic like Trastuzumab. Similar trends between the online and offline techniques were noticed suggesting no inherent bias in the automated analytical workflow that could be readily used for in-process testing during biologics manufacturing.

The advantages of the proposed workflow include; (1) fast turnaround time between sampling from in-process lines; (2) end-to-end integration that is suitable for a variety of different scales ranging from processing samples lab/bench-scale to large commercial scale manufacturing; (3) the ability to configure for both batch/fed-batch bioprocessing modes as well as continuous perfusion modes of cell culture operations; and finally (4) PAT integration to enable rapid process control to meet regulatory requirements of critical quality attributes such as N-glycosylation. One major advantage of the proposal N-GLYcanyzer workflow is the potentially decreased turnaround time between sample removal, rapid N-glycan release using engineered PNGase enzymes, and use of advanced instant glycan labeling chemistry that can further decrease the total sample preparation time spent for mAb protein denaturing and enzymatic deglycosylation followed by instant-fluorescent labeling (e.g., Agilent Instant PC-tag versus classical 2-AB tag) from 3 hours down to less than one hour processing time (data not shown). Future work includes further optimization of the proposed N-GLYcanyzer online platform to rapidly monitor biologics N-glycosylation to enable PAT enabled process control by manipulating upstream cell culture process variables to increase glycosylation quality to within regulatory requirements.

## Acknowledgements

This work was supported by the U. S. Food and Drug Administration (FDA) through the FDA-CBER Award 1R01FD006588. The authors would like to thank the Rutgers FDA project team (Viki Chopda, Haoran Zhang, Doug Hausner, and Antash Chaturvedi), Eppendorf Inc. (Richard Mirro), Agilent Technologies Inc. (Wayne Heacock and Oscar Potter), and FIALabs (Dan Hasle, Scott Schroeder, and Ilkka Lahdesmaki) for their extensive and timely support of this project. Special thanks to Genscript Biotech Corporation (Piscataway, NJ) for the engineered CHO cell line gift to Rutgers University.

## References

1. Coffman, J. et al. A common framework for integrated and continuous biomanufacturing. Biotechnol. Bioeng. 118, 1721–1735 (2021).

2. Konstantinov, K. B. & Cooney, C. L. White paper on continuous bioprocessing May 20-21, 2014 continuous manufacturing symposium. J. Pharm. Sci. 104, 813–820 (2015).

3. Fisher, A. C. et al. The Current Scientific and Regulatory Landscape in Advancing Integrated Continuous Biopharmaceutical Manufacturing. Trends Biotechnol. 37, 253–267 (2019).

4. Chopda, V. et al. Recent advances in integrated process analytical techniques, modeling, and control strategies to enable continuous biomanufacturing of monoclonal antibodies. J. Chem. Technol. Biotechnol. jctb.6765 (2021). doi:10.1002/jctb.6765

5. Wasalathanthri, D. P. et al. Technology outlook for real-time quality attribute and process parameter monitoring in biopharmaceutical development—A review. Biotechnol. Bioeng. 117, 3182–3198 (2020).

6. Pollock, J., Coffman, J., Ho, S. V & Farid, S. S. Integrated continuous bioprocessing: Economic, operational, and environmental feasibility for clinical and commercial antibody manufacture. Biotechnol. Prog. 33, 854–866 (2017).

7. Guidance for Industry PAT — A Framework for Pharmaceutical Development, Manufacturing, and Quality Assurance. (2004).

8. Kelley, B. Industrialization of mAb production technology: The bioprocessing industry at a crossroads. MAbs 1, 443–452 (2009).

9. Alt, N. et al. Determination of critical quality attributes for monoclonal antibodies using quality by design principles. Biologicals 44, 291–305 (2016).

10. Liu, L. Antibody glycosylation and its impact on the pharmacokinetics and pharmacodynamics of monoclonal antibodies and Fc-fusion proteins. J. Pharm. Sci. 104, 1866–1884 (2015).

11. Carrillo-Cocom, L. M. et al. Amino acid consumption in naïve and recombinant CHO cell cultures: producers of a monoclonal antibody. Cytotechnology 67, 809–820 (2015).

12. Fan, Y. et al. Amino acid and glucose metabolism in fed-batch CHO cell culture affects antibody production and glycosylation. Biotechnol. Bioeng. 112, 521–535 (2015).

13. Traustason, B. Amino Acid Requirements of the Chinese Hamster Ovary Cell Metabolism during Recombinant Protein Production. 1–22 (2019). doi:10.1101/796490

14. Ivarsson, M., Villiger, T. K., Morbidelli, M. & Soos, M. Evaluating the impact of cell culture process parameters on monoclonal antibody N-glycosylation. J. Biotechnol. 188, 88–96 (2014).

15. Kimzey, M. et al. Development of an Instant Glycan Labeling Dye for High Throughput Analysis by Mass Spectrometry. ProZyme 25, 1295–1295 (2015).

16. Lauber, M. A. et al. Rapid Preparation of Released N -Glycans for HILIC Analysis Using a Labeling Reagent that Facilitates Sensitive Fluorescence and ESI-MS Detection. Anal. Chem. 87, 5401–5409 (2015).

17. Martín, M. A. et al. A novel and rapid strategy for labelling N -glycans and glycopeptides for qualitative analysis using UHPLC-ESI-MS. 765502

18. Chu, A. H. A., Saati, A. E., Scarcelli, J. J., Cornell, R. J. & Porter, T. J. Reactivity-driven cleanup of 2-Aminobenzamide derivatized oligosaccharides. Anal. Biochem. (2018). doi:10.1016/j.ab.2018.01.012

19. Wu, M. et al. An ultrafast and highly efficient enrichment method for both N-Glycopeptides and N-Glycans by bacterial cellulose. Anal. Chim. Acta 1140, 60–68 (2020).

20. Sha, S., Handelman, G., Liu, N., Xie, D. & Yoon, S. At-line N-linked glycan profiling for monoclonal antibodies with advanced sample preparation and high-performance liquid chromatography. J. Biosci. Bioeng. 130, 327–333 (2020).

21. Tharmalingam, T., Wu, C. H., Callahan, S. & Goudar, C. T. A framework for real-time glycosylation monitoring (RT-GM) in mammalian cell culture. Biotechnol. Bioeng. (2015). doi:10.1002/bit.25520

22. Liu, Y. et al. A Fully Integrated Online Platform For Real Time Monitoring Of Multiple Product Quality Attributes In Biopharmaceutical Processes For Monoclonal Antibody Therapeutics. J. Pharm. Sci. (2021). doi:10.1016/j.xphs.2021.09.011

23. Kozak, R. P., Tortosa, C. B., Fernandes, D. L. & Spencer, D. I. R. Comparison of procainamide and 2-aminobenzamide labeling for profiling and identification of glycans liquid chromatography with fluorescence detection coupled to electrospray ionization-mass spectrometry. Anal. Biochem. 486, 38–40 (2015).

24. Ruhaak, L. R., Steenvoorden, E., Koeleman, C. A. M., Deelder, A. M. & Wuhrer, M. 2-Picoline-borane: A non-toxic reducing agent for oligosaccharide labeling by reductive amination. Proteomics 10, 2330–2336 (2010).

25. Cordes, E. H. & Jencks, W. P. On the Mechanism of Schiff Base Formation and Hydrolysis. J. Am. Chem. Soc. 84, 832–837 (1962).

26. Fang, J., Qin, G., Ma, J. & She, Y. M. Quantification of plant cell wall monosaccharides by reversed-phase liquid chromatography with 2-aminobenzamide pre-column derivatization and a non-toxic reducing reagent 2-picoline borane. J. Chromatogr. A 1414, 122–128 (2015).

27. Vreeker, G. C. M. & Wuhrer, M. Reversed-phase separation methods for glycan analysis. Anal. Bioanal. Chem. 409, 359–378 (2017).

28. Anumula, K. R. Single tag for total carbohydrate analysis. Anal. Biochem. 457, 31–37 (2014).

29. Orrego, A. H. et al. Stabilization of enzymes by multipoint covalent attachment on aldehyde-supports: 2-picoline borane as an alternative reducing agent. Catalysts 8, (2018).

30. Melmer, M., Stangler, T., Premstaller, A. & Lindner, W. Comparison of hydrophilic-interaction, reversed-phase and porous graphitic carbon chromatography for glycan analysis. J. Chromatogr. A 1218, 118–123 (2011).

31. Mauko, L. et al. Comparison of ZIC-HILIC and graphitized carbon-based analytical approaches combined with exoglycosidase digestions for analysis of glycans from monoclonal antibodies. Journal of Chromatography B: Analytical Technologies in the Biomedical and Life Sciences 911, 93–104 (2012).

32. Ashwood, C., Pratt, B., Maclean, B. X., Gundry, R. L. & Packer, N. H. Standardization of PGC-LC-MS-based glycomics for sample specific glycotyping. Analyst 144, 3601–3612 (2019).

33. Wei, J. et al. Toward Automatic and Comprehensive Glycan Characterization by Online PGC-LC-EED MS/MS. Anal. Chem. 92, 782–791 (2020).

34. Melmer, M. Investigation of High-Performance Liquid Chromatography Methods for the Analysis of Protein N-glycans. Naturwissenschaften (University of Vienna, 2010).

35. Madabhushi, S. R., Podtelezhnikov, A. A., Murgolo, N., Xu, S. & Lin, H. Understanding the effect of increased cell specific productivity on galactosylation of monoclonal antibodies produced using Chinese hamster ovary cells. J. Biotechnol. 329, 92–103 (2021).

