## Supplemental Information for "An Integrated Process Analytical Platform for Automated Monitoring of Monoclonal Antibody N-linked Glycosylation"

Title:

List of Authors:

Aron Gyorgypal[1], and Shishir P.S. Chundawat[1,*]

Affiliation:

[1]Department of Chemical and Biochemical Engineering, Rutgers University, Piscataway, New Jersey, United States

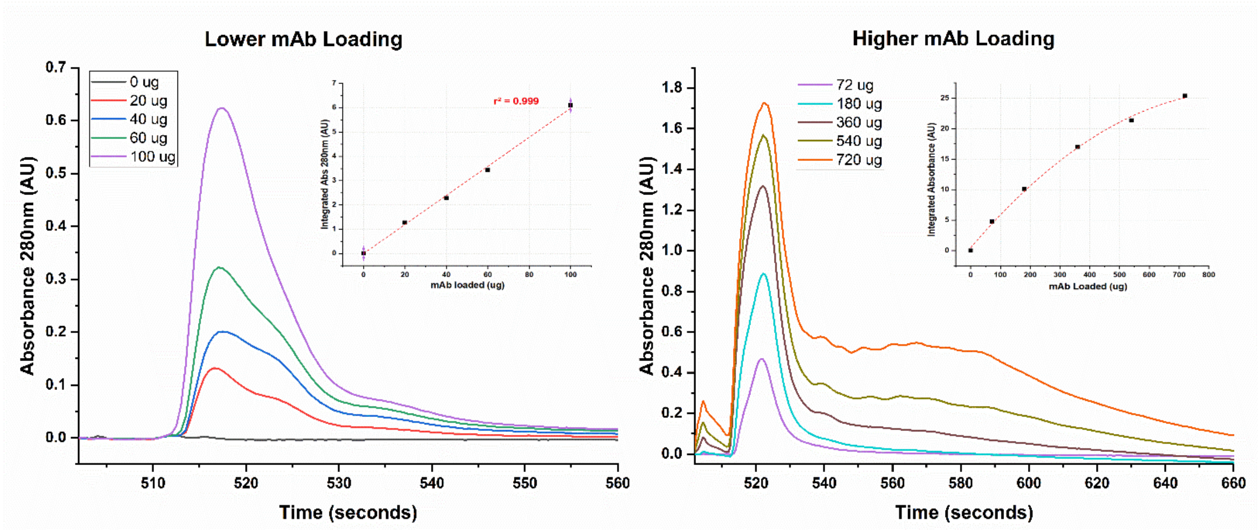


**Supplemental Figure 1 (S1). Protein A Loading Optimization:** Loading curves for a 30 mm length packed protein A column was conducted. Here, S1A shows the elution profile for mAb at varying loading concentrations (0, 20, 40, 60, 100 μg) of mAb loaded on to the column. Under a 20 second integration the loading curve shows a strong linear correlation for the mAb loading of $r^2$ =0.999. A further increase in mAb loading on the calibration curve is shown in S1B. Briefly, tailing can be seen for the peaks upon increasing the mAb loading. This can be attributed to the weak elution strength of 0.1% formic acid. The use of an ionic modifier (salt) could help mitigate the tailing issue but would also increase the total salt concentration in the eluted sample.

| Oligomer | Reducer | Aqueous 2AB | ± | | Aqueous 2AA | ± | | Non-Aqueous 2AB | ± | | Non-Aqueous 2AA | ± | |
|---|---|---|---|---|---|---|---|---|---|---|---|---|---|
| **Glc-4** | **2-Picoline Borane** | 96% | ± | 5% | 2% | ± | 1% | 8% | ± | 0% | 2% | ± | 3% |
| | **Sodium Cyanoborohydride** | 100% | ± | 3% | 10% | ± | 0% | 97% | ± | 3% | 15% | ± | 1% |
| | **No Reducer** | 17% | ± | 7% | 0% | ± | 1% | 22% | ± | 0% | 0% | ± | 0% |
| **Glc-5** | **2-Picoline Borane** | 97% | ± | 6% | 14% | ± | 1% | 17% | ± | 1% | 11% | ± | 3% |
| | **Sodium Cyanoborohydride** | 100% | ± | 2% | 13% | ± | 1% | 99% | ± | 1% | 22% | ± | 1% |
| | **No Reducer** | 15% | ± | 5% | 1% | ± | 1% | 15% | ± | 0% | 0% | ± | 0% |
| **Glc-6** | **2-Picoline Borane** | 98% | ± | 10% | 17% | ± | 0% | 19% | ± | 2% | 17% | ± | 2% |
| | **Sodium Cyanoborohydride** | 100% | ± | 3% | 15% | ± | 1% | 99% | ± | 2% | 22% | ± | 4% |
| | **No Reducer** | 17% | ± | 1% | 1% | ± | 0% | 12% | ± | 0% | 0% | ± | 0% |
| **Glc-7** | **2-Picoline Borane** | 100% | ± | 8% | 22% | ± | 1% | 15% | ± | 1% | 17% | ± | 1% |
| | **Sodium Cyanoborohydride** | 91% | ± | 7% | 17% | ± | 1% | 84% | ± | 1% | 25% | ± | 2% |
| | **No Reducer** | 14% | ± | 4% | 1% | ± | 1% | 9% | ± | 0% | 0% | ± | 0% |
| **Glc-8** | **2-Picoline Borane** | 100% | ± | 8% | 31% | ± | 1% | 12% | ± | 0% | 25% | ± | 0% |
| | **Sodium Cyanoborohydride** | 83% | ± | 5% | 25% | ± | 0% | 68% | ± | 1% | 34% | ± | 2% |
| | **No Reducer** | 13% | ± | 4% | 2% | ± | 1% | 7% | ± | 0% | 2% | ± | 0% |
| **Glc-9** | **2-Picoline Borane** | 100% | ± | 2% | 37% | ± | 0% | 6% | ± | 1% | 24% | ± | 0% |
| | **Sodium Cyanoborohydride** | 21% | ± | 0% | 23% | ± | 0% | 22% | ± | 1% | 33% | ± | 2% |
| | **No Reducer** | 9% | ± | 4% | 2% | ± | 1% | 4% | ± | 1% | 0% | ± | 0% |
| **Glc-10** | **2-Picoline Borane** | 100% | ± | 1% | 47% | ± | 0% | 5% | ± | 2% | 25% | ± | 3% |
| | **Sodium Cyanoborohydride** | 18% | ± | 0% | 23% | ± | 0% | 16% | ± | 0% | 34% | ± | 1% |
| | **No Reducer** | 7% | ± | 3% | 1% | ± | 1% | 2% | ± | 0% | 1% | ± | 0% |
| **Glc-11** | **2-Picoline Borane** | 100% | ± | 7% | 78% | ± | 0% | 7% | ± | 2% | 31% | ± | 5% |
| | **Sodium Cyanoborohydride** | 16% | ± | 0% | 13% | ± | 1% | 17% | ± | 5% | 51% | ± | 2% |
| | **No Reducer** | 6% | ± | 0% | 4% | ± | 0% | 4% | ± | 2% | 0% | ± | 0% |
| **Glc-12** | **2-Picoline Borane** | 100% | ± | 3% | 99% | ± | 1% | 7% | ± | 6% | 36% | ± | 5% |
| | **Sodium Cyanoborohydride** | 23% | ± | 2% | 25% | ± | 3% | 21% | ± | 7% | 54% | ± | 8% |
| | **No Reducer** | 8% | ± | 1% | 7% | ± | 1% | 8% | ± | 2% | 7% | ± | 1% |

**Supplemental Figure 2 (S2). Dextran labeling efficiency depends on reducing agent:** Labeling efficiencies can be seen in this chart for a dextran ladder ranging from a 4-mer to 12-mer malto-oligomers. The use of 2-picoline borane under aqueous conditions for all oligomers gave the highest level of labeling efficiency while the non-aqueous labeling conditions always gave lower efficiency. For all oligomers, the use of 2-picoline borane with 2-AB dye seemed to give the best efficiencies in glycan labeling.

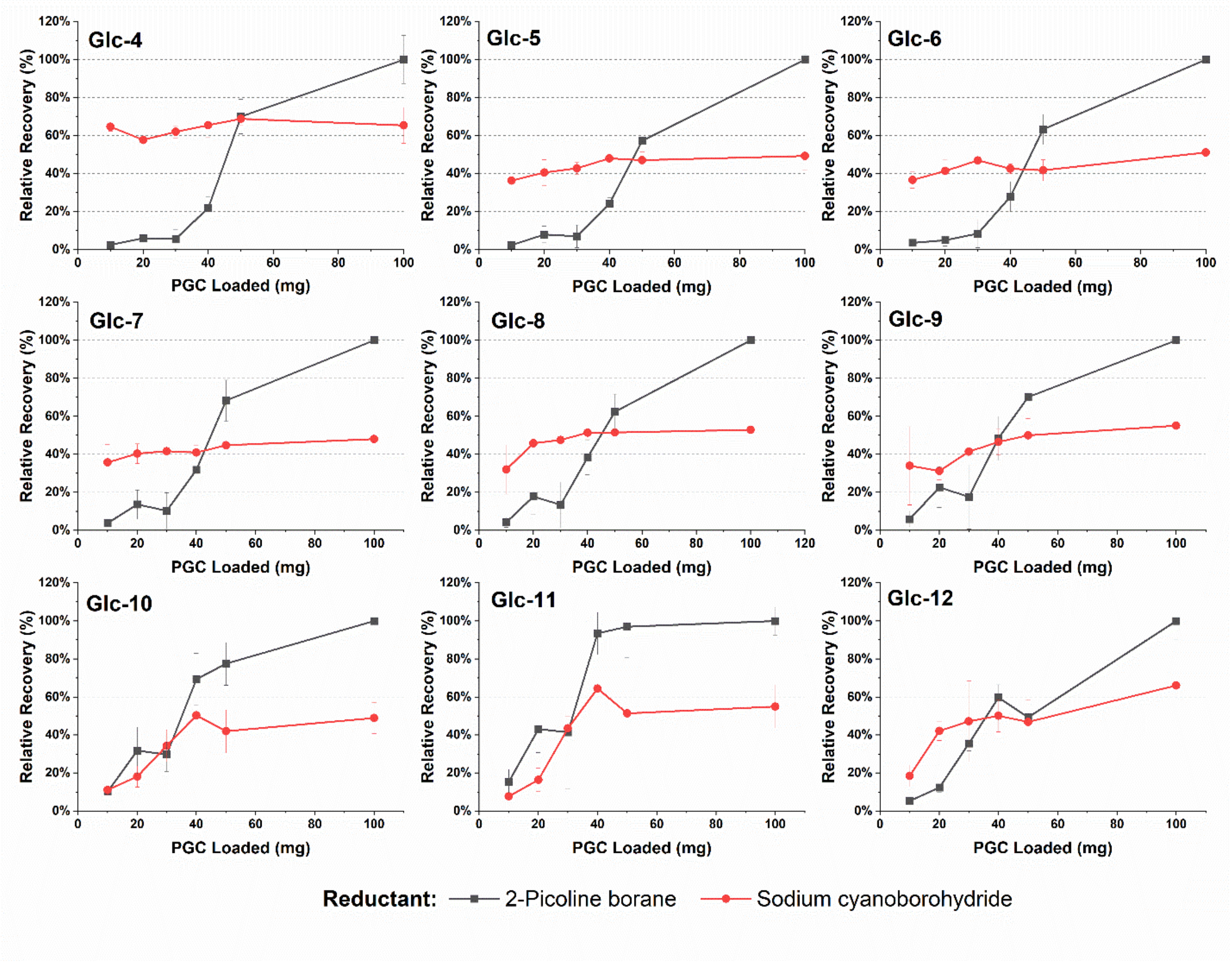


**Supplemental Figure 3 (S3). Impact of PGC bed weight on labeled dextran ladder recovery:** Recovery Rates for 2-AB labeled dextran ladder ranging between 4 to 12 glucose units was determined. With an increase in PGC bed weight an increase in 2-picoline borane reduced 2-AB labeled oligomers can be seen. However, a constant recovery of 2-AB labeled glycans with marginal increase is seen when using sodium cyanoborohydride as the reductant.
